# Tracking bilingual brain processing at high spatiotemporal resolution reveals where and when second language slows down

**DOI:** 10.64898/2026.02.01.701554

**Authors:** Teng Ieng Leong, Victoria Lai Cheng Lei, Jian Hwee Ang, Cheok Teng Leong, Chi Un Choi, Martin I. Sereno, Defeng Li, Ruey-Song Huang

**Affiliations:** Centre for Cognitive and Brain Sciences, University of Macau, Taipa, Macau SAR, China; Faculty of Arts and Humanities, University of Macau, Taipa, Macau SAR, China; Faculty of Science and Technology, University of Macau, Taipa, Macau SAR, China; Department of Cognitive Science, University of California, San Diego, La Jolla, CA, USA

**Keywords:** phase-encoded fMRI, traveling waves, language comprehension, overt speech production, dual-stream models

## Abstract

A central question in bilingualism research is whether challenges in a second language (L2) reflect a fundamentally different processing route from the first language (L1), or slower processing at particular stages. Although functional magnetic resonance imaging (fMRI) revealed substantial cortical overlap between L1 and L2, different processing mechanisms may interleave within the overlap. Conventional fMRI methods lack the combined spatial and temporal resolution needed to determine precisely where and when L1 and L2 dissociations occur. Here, we used rapid phase-encoded fMRI to track L1 and L2 processing within overlapping cortical regions. Thirty-one Chinese-English bilinguals completed sentence-level reading, listening, reading-aloud, and shadowing tasks in both Chinese (L1) and English (L2). By analyzing hemodynamic traveling waves along sample paths through the visual, auditory, and motor systems, we identified temporal dissociations between L1 and L2 processing. Compared with L1, L2 exhibited significant and consistent delays along dorsal and ventral visual streams during reading and reading aloud. Along auditory streams, L2 delays differed between externally presented speech (listening) and self-generated speech (reading aloud). In the frontal cortex, reading aloud and shadowing exhibited similar progression patterns, but with different temporal offsets between L1 and L2 in the dorsal and ventral motor streams. These findings reveal task- and modality-dependent differences between L1 and L2 processing, and pinpoint the exact locations for delayed L2 processing across multimodal streams. This study suggests that L2 disadvantage is often a matter of timing and efficiency within largely shared brain networks, rather than due to the use of different brain regions.

**Significant Statement:** Rapid phase-encoded fMRI introduces a temporal dimension for comparing L1 and L2 processing, tracking the precise timing of neural information flow across the brain. Moving beyond functional localization, we directly visualize and quantify the propagation of hemodynamic traveling waves along selected paths in visual, auditory, and motor systems. Our ability to characterize temporal delays along specific functional streams represents an advancement in understanding dynamic cognitive processes beyond language. The findings can influence theories of language acquisition and cognitive processing in bilingual individuals. The methods to pinpoint locations where delays occur can be applied to other areas of cognitive neuroscience, allowing the exploration of complex neural interaction in real time across sensory, motor, and cognitive domains.

## Introduction

While the challenges associated with learning or using a second language are widely acknowledged, the precise neural dynamics that differentiate the processing of a second language from a first remain to be fully elucidated. Previous research indicates that cortical representations of first and second languages (L1 and L2) overlap to a considerable extent in bilinguals (Indefrey, 2006; Abutalebi, 2008; Liu and Cao, 2016; Cargnelutti et al., 2019; Sulpizio et al., 2020; Malik-Moraleda et al., 2024). However, overlap in brain activation regions does not necessarily imply shared underlying neural mechanisms, as distinct neural populations and computations for processing different languages could be interleaved within the same region (Hernandez et al., 2005; Xu et al., 2017). Beyond reporting the differences in activation centers, extent, strength, and patterns, previous contrast-based neuroimaging studies lacked additional measures to compare how different languages are processed in the brain (Chee et al., 1999; Fabbro, 2001; Buchweitz et al., 2012; Kim et al., 2016). More importantly, language processing is a dynamic process that involves the simultaneous engagement of parallel and sequential streams (Price, 2012). For reading, a dorsal visual stream is proposed for sub-lexical processing and a ventral visual stream for lexical analysis (Pugh et al., 2000; Borowsky et al., 2006; Sood and Sereno, 2016). Similarly, for speech perception, a dorsal auditory stream maps sound to articulation, while a ventral auditory stream maps sound to meaning (Hickok and Poeppel, 2007; Saur et al., 2008; Rauschecker and Scott, 2009). A comparable dual-system exists for speech production, with dorsal premotor areas coordinating pitch and ventral premotor areas coordinating phonemic selection (Hickok et al., 2023). Therefore, comparing how different languages are processed dynamically in the brain will require investigating information flows through these multimodal streams.

However, uncovering where and when the differences between languages occur involves continuous tracking of brain activities across regions, which poses methodological challenges. While intracranial recordings can detect spatiotemporal brain dynamics with millimeter and millisecond resolutions, the coverage of cortical regions is limited by the placement of electrodes (Woolnough et al., 2019; Castellucci et al., 2022; Leonard et al., 2024; Kovacs et al., 2025; Norman-Haignere et al., 2025). This affects the ability to capture a complete picture of information flows across distant brain regions simultaneously (Parvizi and Kastner, 2018). Noninvasive neuroimaging techniques, such as electroencephalography (EEG), are highly sensitive to artifacts during speech production and cannot precisely track brain activities due to spatial smearing (Burle et al., 2015). While fMRI offers excellent spatial resolution, it has long been thought that the slow hemodynamic response and low sampling rate (e.g., 1 s per volume) cannot capture the temporal dynamics of language processing unfolding over tens to hundreds of milliseconds (Friederici, 2017). Nevertheless, recent fMRI studies have successfully unravelled the temporal sequence of activations across brain regions during visually guided actions and natural language processing tasks (Chen et al., 2019; Lei et al., 2024). By adding a temporal dimension to the static activation map, rapid phase-encoded fMRI can track down the direction and timing of neural information flow via streams of hemodynamic traveling waves over the cortical surface (Sereno et al., 2022; Lei et al., 2024).

Here, thirty-one sequential bilinguals participated in a rapid phase-encoded fMRI experiment, in which they performed four sentence-level language perception and production tasks in their first and second languages. The phases of periodic hemodynamic fluctuations at the task frequency were displayed on the cortical surface, revealing activations with different phases across the brain. Based on dual-stream models (Pugh et al., 2000; Borowsky et al., 2006; Hickok and Poeppel, 2007; Saur et al., 2008; Rauschecker and Scott, 2009; Hickok et al., 2023), we sampled selected paths through surface traveling waves propagating across shared cortical regions. We then compared the activation phases between L1 and L2 along each path in the visual, auditory, and motor systems. The goal was to pinpoint precise locations of temporal dissociations between L1 and L2 processing along these paths.

## Materials and Methods

### Participants

The study analyzed fMRI datasets of 31 subjects (22 females, 9 males; mean age: 23.3 ± 4 years) recruited by the Brain and Language Project at the University of Macau (Lei et al., 2024). All subjects were sequential bilinguals, with Chinese (Mandarin) as their first language and English as their second language, which they acquired between the ages of 4 and 13 (mean age of acquisition: 7.1± 2.1 years). The subjects’ English proficiency was assessed using their most recent standardized English language test (IELTS 6.5+ or equivalent). All subjects had normal or corrected-to-normal vision and no history of neurological impairment. Written informed consent was obtained from all subjects in accordance with the experimental protocols approved by the Ethics Committee of the University of Macau.

### Stimulus construction and presentation

Language stimuli consisted of 128 simple sentences in Chinese and 128 in English (Lei et al., 2024). Chinese sentences were systematically constructed with 14 characters using words selected from the Lancaster Corpus of Mandarin Chinese (https://www.lancaster.ac.uk/fass/projects/corpus/LCMC/). English sentences contained 15–17 syllables and were built using high-frequency verbs from the Longman Communication 3000 list, with vocabulary verified to be of mid to high frequency. All sentences in both languages were constructed in a subject-verb-object (S-V-O) structure and validated by a linguistics professor to ensure they are plausible, unambiguous, and grammatically natural. Visual stimuli were presented in black text centered on a white background on an LCD monitor (InroomViewingDevice, NordicNeuroLab AS). Auditory stimuli were synthesized using the Google Cloud Text-to-Speech tool under default settings and delivered via MR-compatible headphones (OptoActive II, OptoAcoustics Ltd.). An MR-compatible microphone (OptoAcoustics Ltd.) was used to record overt speech output while also providing real-time auditory feedback through the headphones.

### Experimental design

The current study included the following language processing tasks in one Chinese session and one English session, respectively: (i) Reading: subjects silently read and comprehended a written sentence without any response or engagement; (ii) Listening: subjects attentively listened to and comprehended a spoken sentence without any response or engagement; (iii) Reading aloud: subjects read a written sentence aloud; (iv) Shadowing: subjects listened to and repeated a spoken sentence simultaneously. Each task was performed in two nonconsecutive 256-s scans, each consisting of sixteen 16-s trials. There were two phases in each trial, which occurred periodically in 256 s. In phase 1 (task period), a sentence stimulus was presented either visually (for reading and reading aloud) or auditorily (for listening and shadowing) within the first 5 s of each 16-s trial. In phase 2 (rest period), a blank white screen was presented for the remaining 11 s of the trial.

### Speech recording and analysis

Auditory stimuli and speech output were recorded at 16 KHz using OptiMRI 3.1 software (OptoAcoustics Ltd.), in sync with Transistor-Transistor Logic (TTL) signals from the MRI system. The onset and offset times of speech output in each trial of the reading-aloud and shadowing tasks were identified through manual annotation of the audio recordings using Audacity software (https://www.audacityteam.org).

### Image acquisition

Functional and structural brain images were acquired using a 32-channel head coil in a Siemens MAGNETOM Prisma 3T MRI scanner at the Centre for Cognitive and Brain Sciences, University of Macau. Twelve functional scans were acquired using a blipped-CAIPIRINHA simultaneous multi-slice (SMS), single-shot echo planar imaging (EPI) sequence (acceleration factor: 5; interleaved ascending slices; TR: 1000 ms; TE: 30 ms; flip angle: 60º; 55 axial slices; field of view: 192×192 mm; matrix size: 64×64; voxel size: 3×3×3 mm; bandwidth: 2368 Hz/Px; 256 TRs per image; dummy: 6 TRs; effective scan time: 256 s). Two sets of T1-weighted structural images were acquired using an MPRAGE sequence (TR: 2300 ms; TE: 2.26 ms; TI: 900 ms; flip angle: 8º; 256 axial slices; field of view: 256×256 mm; matrix size: 256×256; voxel size: 1×1×1 mm; bandwidth: 200 Hz/Px; scan time: 234 s) with the same slice centre and orientation as the functional images.

### Image preprocessing

All raw images (DICOM *.ima files) were converted to Analysis of Functional NeuroImages (AFNI; https://afni.nimh.nih.gov) BRIK files using the AFNI *to3d* command. Functional images were motion-corrected by registering all volumes in 12 scans to the first volume (reference) of the seventh scan using the AFNI *3dvolreg* command. Subsequently, the motion-corrected functional images were coregistered to the structural images using the *csurf* package (https://pages.ucsd.edu/~msereno/csurf) (Sereno et al., 2022). For each subject, two T1-weighted structural scans were averaged prior to cortical surface reconstruction using FreeSurfer 7.2 (https://surfer.nmr.mgh.harvard.edu) (Dale, 1999; Fischl et al., 1999).

### Functional data analysis

Functional images underwent a data processing pipeline consisting of voxel-wise Fourier-based analyses and surface-based (vertex-wise) group-averaging methods developed by Sereno and colleagues (Sereno et al., 1995, 2001, 2022; Hagler et al., 2006, 2007). The following analyses were carried out using standard functions included in the *csurf* package (Sereno et al., 2022). In each functional scan (64×64×55×256 sample points), the time series *x*_*m*_*(t)* of voxel *m* was analyzed with a 256-point discrete Fourier transform (Huang et al., 2012; Chen et al., 2017, 2019):

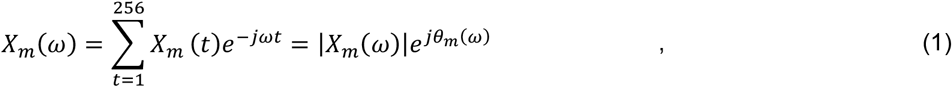

where *X*_*m*_(*ω*) is the Fourier component at frequency *ω* between 0-127 cycles per scan, and |*X*_*m*_(*ω*)| and *θ*_*m*_(*ω*) are its amplitude and phase. The task frequency is defined as (16 cycles per scan), at which the BOLD signals fluctuate periodically in response to periodic stimuli and tasks. The complex signal (*X*_*m*_(*ω*_*s*_)^*R*^, *X*_*m*_(*ω*_*s*_)^*I*^) and noise (*X*_*m*_(*ω*_*n*_)^*R*^, *X*_*m*_(*ω*_*n*_)^*I*^) in voxel *m* are defined as the Fourier components at task frequency *ω*_*s*_ and other non-task frequencies at *ω*_*n*_, respectively. The statistical significance of the signal-to-noise ratio (SNR) in voxel *m* was evaluated by:

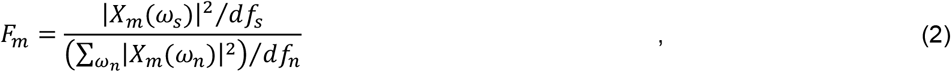

where *df*_*s*_ = 2 (degrees of freedom of real and imaginary components at *ω*_*s*_ = 16 cycles per scan) and *df*_*n*_ = 230 (degrees of freedom of real and imaginary components at non-task frequencies *ω*_*n*_ = excluding 0-3, 15, 17, 31-33, 47-49, and 64 from [0, 127] cycles per scan, i.e., *df*_*n*_ = (128 −4 −2 −3 −3 −1)*2 = 230). The *P*-value of the *F*-ratio (SNR) in Equation 2 was estimated by the cumulative distribution function *F*(*F*_*m*_; *df*_*s*_, *df*_*n*_) (Huang et al., 2012; Chen et al., 2017, 2019). A complex *F*-value, 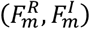, incorporating both the SNR and phase, *θ*_*m*_(*ω*_*s*_), of the periodic signal in each voxel was obtained by:

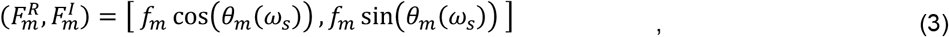

where *f*_*m*_ is the square root of *F*_*m*_. Slice-timing correction was performed by the *csurf fourier* command, which adjusted the computed phase using slice times extracted from the *.ima files (implemented by the AFNI *to3d* command with the “-time” option). For each subject *S* = {1, 2, …, *N*}, the complex *F*-values in corresponding voxels *m* in scan *k* ={1, 2} were vector-averaged across corresponding voxels in two scans for the same language task by:

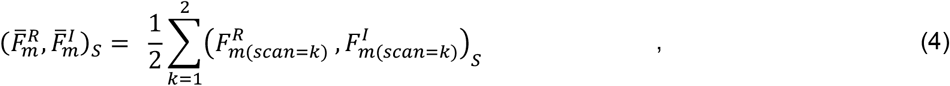

using the “Combine 3D Phase Statistics” function in *csurf*. The resulting subject-level voxel-wise average *F*-values, 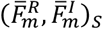, were then projected onto vertex *v* on individualized cortical surfaces of subject *S*, yielding a surface-based map, 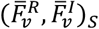, for each task. The phase at vertex *v* in the average map of subject *S* was obtained by:

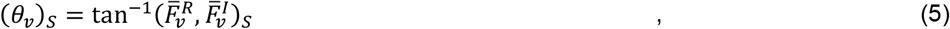

The spherical averaging method (Fischl et al., 1999; Hagler et al., 2006; Sood and Sereno, 2016; implemented as the “Cross Session Spherical Average” function in *csurf*) was used to obtain a surface-based group-average map for each task. First, each surface-based map of subject *S*, 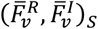, was morphed and resampled to a common spherical coordinate system using the FreeSurfer *mri_surf2surf* command (https://freesurfer.net/fswiki/mri_surf2surf), yielding a new map displayed on the *fsaverage* surface, 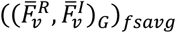. Second, the complex *F*-values at vertex *v* on the *fsaverage* surface were vector-averaged (vertex-wise) across subjects (*N* = 31) by:

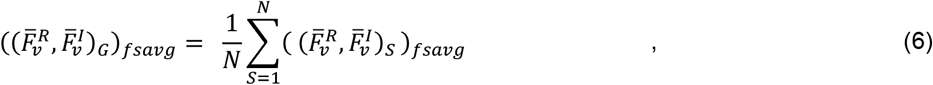

yielding a surface-based group-average map for each task. The amplitude and phase at vertex *v* in this map were obtained by:

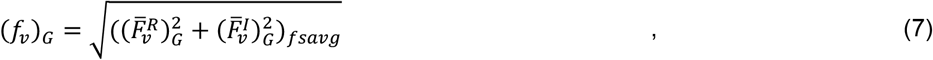

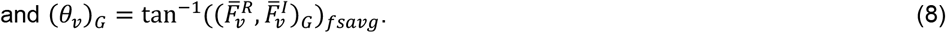

The amplitudes at each vertex in the common spherical coordinate system were tested across subjects by *F*-statistics (*N* = 31, *F*(2, 60) > 4.97, *P* < 0.01), and corrected for multiple comparisons using a surface-based cluster-size exclusion method (Hagler et al., 2006, 2007), with cluster size = 64 mm^2^, alpha = 0.05, *P* < 0.01, cluster-corrected. The resulting phase-encoded group-average activation maps, consisting of amplitude and phase at each vertex (Equations 7 and 8) were displayed on the inflated and flattened cortical surfaces of *fsaverage* using the *csurf* package (Fig. 1*A, B, C, D*). The group-average activation phase, (*θ*_*v*_)_*G*_, ranging from 0 to 2*π* radians) at vertex *v* was converted to a time delay (ranging from 0 to 16 s). In the current study, we only analyzed and displayed task-positive activations with phases between 3 and 10 s, accounting for the hemodynamic response delay. Task-negative activations, such as those in the default mode network, exhibiting phases beyond this range were not shown in the current study and will be reported elsewhere.

**Figure 1.**
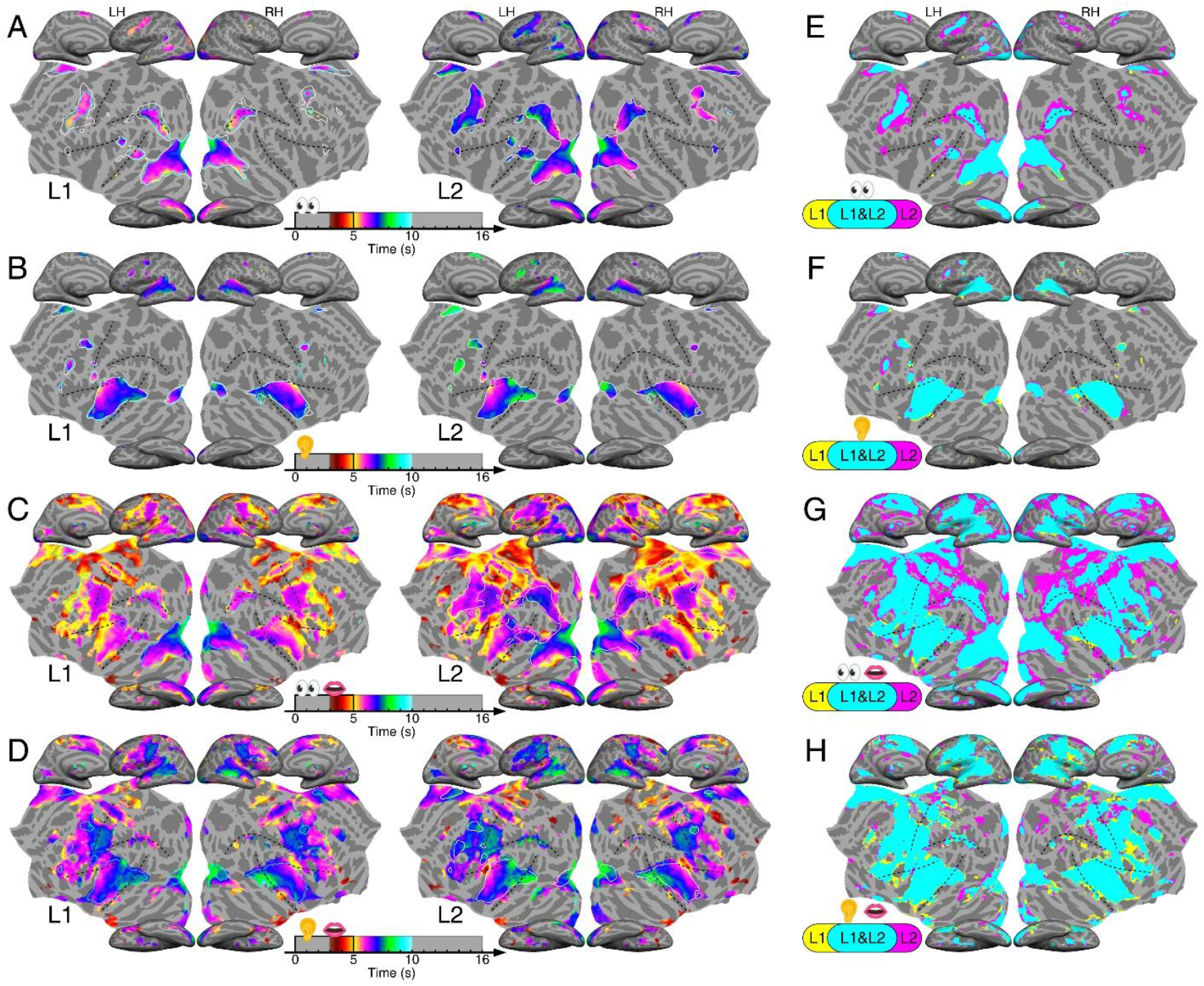
Phase-encoded activation maps for four tasks in L1 and L2 and L1-L2 conjunction maps. ***A*** to ***D***, Group-average maps showing regions with significant periodic activations (*N* = 31, *P* < 0.01, cluster-corrected) during reading (***A***), listening (***B***), reading aloud (***C***), and shadowing (***D***). First column: Chinese (L1), second column: English (L2). LH and RH: left and right hemispheres. Dashed black curves mark the central sulcus, lateral sulcus (Sylvian fissure), STS, and IPS. The phases (time delays) of task-related activations are color-coded (brown = earliest; cyan = latest). For reading (***A***) and listening (***B***), activation contours are shown in cyan (L1) and white (L2). For reading aloud (***C***) and shadowing (***D***), activation maps are overlaid with contours from (***A***) and (***B***) tasks in the same language, respectively. ***E*** to ***H***, conjunction maps for L1-L2 maps in ***A*** to ***D***. Yellow, L1-only activations; magenta, L2-only activations; cyan, regions activated by both languages.

### Conjunction analysis

A surface-based conjunction map (Fig. 1*E, F, G, H*) was created for comparing the activation extent between paired L1 and L2 group-average phase-encoded activation maps for each task (Fig. 1*A, B, C, D*). At the same statistical threshold (activation SNR: *F*(2, 230) > 4.7, *P* < 0.01; group statistics: *F*(2, 60) > 4.97, *P* < 0.01, cluster corrected), a vertex on the *fsaverage* surface was colored as follows: yellow for significant activation only in the L1 map; magenta for significant activation only in the L2 map; and cyan for significant activation in both L1 and L2 maps (i.e., overlap). The percentages of vertices activated by a single language or both languages are summarized in Table 2.

**Table 1.** Group-average timings of recorded speech output.

| Language | Task | Speech onset (s)<br>Mean ± s.d. | Speech offset (s)<br>Mean ± s.d. | Duration (s)<br>Mean ± s.d. |
| --- | --- | --- | --- | --- |
| Chinese (L1) | Reading aloud | 1.16 ± 0.15 | 4.17 ± 0.37 | 3.03 ± 0.32 |
| | Shadowing | 1.40 ± 0.36 | 5.49 ± 0.36 | 4.09 ± 0.27 |
| English (L2) | Reading aloud | 1.22 ± 0.17 | 5.09 ± 0.47 | 4.06 ± 0.44 |
| | Shadowing | 1.30 ± 0.27 | 5.89 ± 0.48 | 4.60 ± 0.36 |

$N = 31$ ; s.d., standard deviation.

**Table 2.** Percentages of L1, L2, and overlapping regions in the conjunction maps.

| Tasks | Left hemisphere |  |  |  | Right hemisphere |  |  |  |
| --- | --- | --- | --- | --- | --- | --- | --- | --- |
|  | Chinese (L1) |  | English (L2) |  | Chinese (L1) |  | English (L2) |  |
| Reading | 1.3% | 98.7% | 57.6% | 42.4% | 3.1% | 96.9% | 51.9% | 48.1% |
| Listening | 6.7% | 93.3% | 90.1% | 9.9% | 10.0% | 90.0% | 93.4% | 6.6% |
| Reading aloud | 2.0% | 98.0% | 69.9% | 30.1% | 2.7% | 97.3% | 62.3% | 37.7% |
| Shadowing | 11.0% | 89.0% | 87.8% | 12.2% | 15.9% | 84.1% | 82.7% | 17.3% |

### Traveling wave analysis

To compare fine-grained spatiotemporal activation patterns across languages, we first outlined visual, auditory, and motor regions in the flattened cortical surface (as indicated by the boxes in Fig. 2*A*). We then superimposed isophase contours onto the group-average phase-encoded activation maps (Fig. 1), revealing the underlying traveling wave formations in each region for L1 and L2 tasks (Figs. 2-4). This was carried out by setting the parameter $phasecontoursflag to 1 in the tksurfer interface of *csurf*. As an initial step toward comparing processing timings across languages, we sampled multiple paths, each traversing approximately perpendicular to the isophase contours (i.e., ‘wavefronts’) on the cortical surface, based on dual-stream models in visual, auditory, and motor systems (Figs. 2-4) (Hickok and Poeppel, 2007; Saur et al., 2008; Rauschecker and Scott, 2009; Huang and Sereno, 2018; Hickok et al., 2023). To maximize coverage, we delineated each path to span slightly beyond regions exhibiting significant activations in both language maps. The HCP-MMP1 atlas (Glasser et al., 2016) and CsurfMaps1 atlas (Sereno et al., 2022) were used as guide maps for locating these paths on the cortical surface (Fig. 2*B, C*). Table 3 summarizes the number of selected nodes (vertices) on each path and the surface-based regions of interest (sROIs) it passes through. For each path, we computed the mean and standard deviation of phases, (*θ*_*v*_)_*G*_, of 121 vertices (center and 120 neighbors; obtained by 10 steps of smoothing) within an average radius between 3.75 and 5.36 mm centered at each node (vertex) along the path in a group-average activation map. The results were plotted for each task and language (Figs. 2*E*, 2*G*, 3*B*, 3*D*, 3*F*, 4*B*, 4*D*).

**Table 3.** Surface-based regions of interest on visual, auditory, and motor paths according to two cortical parcellation atlases.

| Modality | Path | HCP-MMP1 (Fig. 2B) | CsurfMaps1 (Fig. 2C) | Number of nodes |
| --- | --- | --- | --- | --- |
| Visual | DVS | V3B, IP0, IPS1, MIP, IP1, LIPd, IP2 | V3A, cIPS, LIP0, LIP1, IPS5, N/D | 30 |
|  | VVS | V8, FFC, PH, PHT, TE1p | V8, FFC, PH, N/D | 17 |
| Auditory | Spt | RI, A1, LBelt, PSL, PFcm, PF | CM, N/D, CP, STV2, PSaud1 | 18 |
|  | STS | RI, A1, LBelt, PBelt, A4, A5, TPOJ1, STSdp, STSvp | MM, CM, CL, ML, CP, C_A4, M_A4, N/D | 21 |
|  | ATL | RI, MBelt, A1, PBelt, A4, A5, TA2, STGa | MM, A1, R, RT, TA2, TA3, N/D | 19 |
| Motor | DMS | 6a, FEF, 55b, 4, 3a, 3b | 6a, FEF, PZ-a,v,s, PZ-a,s, 3a_face, 3b_face | 23 |
|  | VMS | IFSa, IFSp, IFJa, IFJp, 6r, 6v, 4, 3a, 3b | N/D, DLPFCa, DLPFC, N/D, 4_face, 3a_face, 3b_face | 25 |
|  | AI | AVI, FOP5, FOP4, FOP1, 43, OP4 | FOPaud, 45aud, N/D, FOP2, N/D, 43aud, S_II | 21 |

N/D, not defined regions.

**Figure 2.**
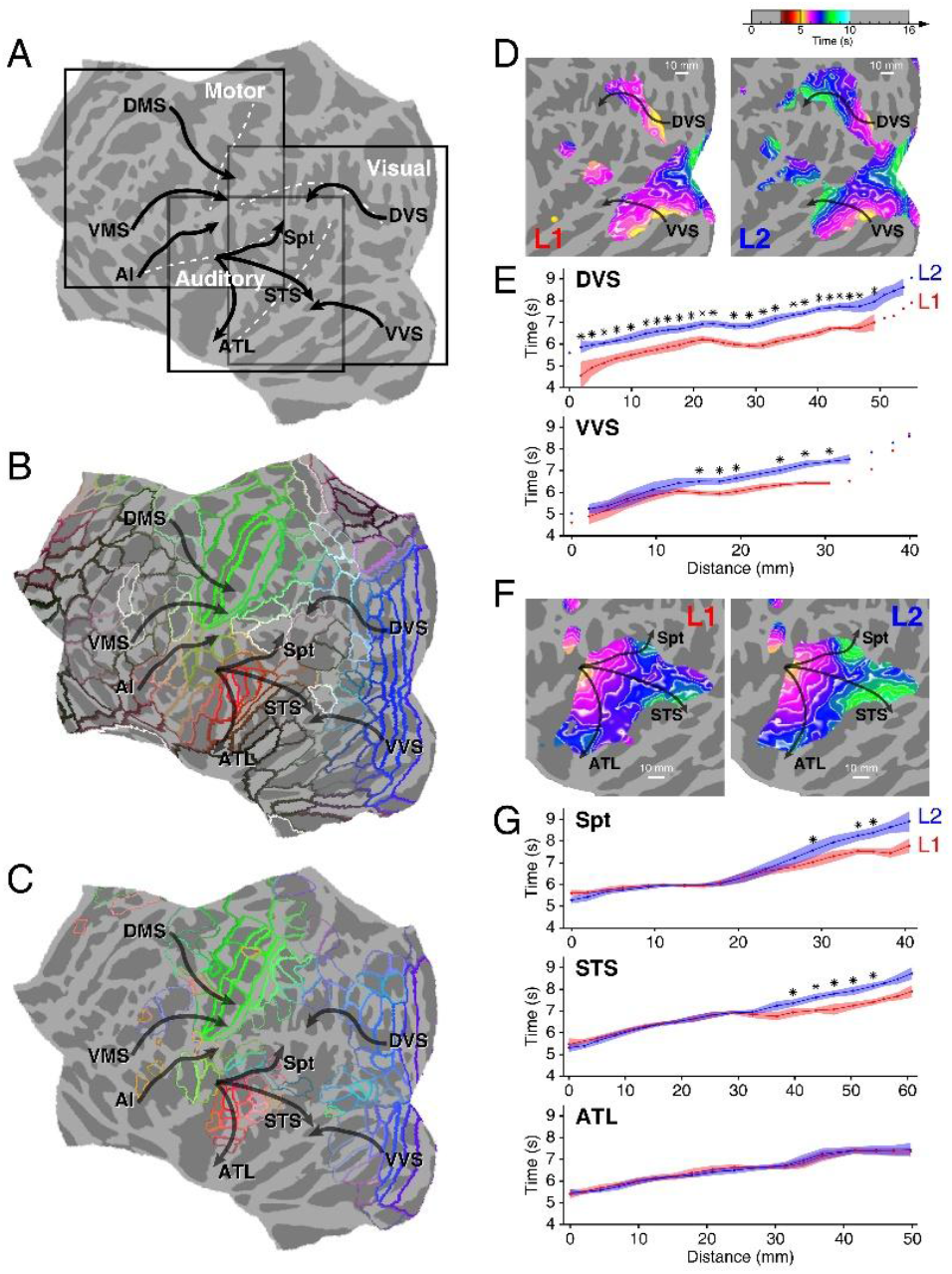
Maps of multimodal paths and analyses of processing times along visual and auditory paths. ***A***, Visual, auditory, and motor paths displayed on the flattened left hemisphere of *fsaverage*. ***B***, Multimodal paths illustrated with the HCP-MMP1 atlas (Glasser et al., 2016). ***C***, Multimodal paths illustrated with the CsufMaps1 topological atlas (Sereno et al., 2022). ***D***, Close-up views of hemodynamic traveling waves propagating through the dorsal visual stream (DVS) and ventral visual stream (VVS) during Chinese (L1; left panel) and English (L2; right panel) reading tasks (see full-hemisphere maps in Fig. 1*A*). Each white contour represents an iso-phase wavefront, indicating vertices with the same activation phase. ***E***, The curve and shading (red: L1; blue: L2) depict the mean and standard deviation of activation phases in 121 vertices (one central vertex and 120 neighbors) surrounding each selected node (vertex) along each path in ***D***. Unconnected dots on some paths indicate nodes with insignificant activations; the paths were selected to cover the larger activation extent of the two language maps. Asterisks (*) indicate significant delays in L2 compared to L1 activation phases across subjects at each node (*P* < 0.05, Watson-Williams test, *N* = 31; see Suppl. Dataset S1 for details). ***F***, Close-up views of hemodynamic traveling waves propagating through the superior temporal cortex during L1 and L2 listening tasks (see full-hemisphere maps in Fig. 1*B*). Three sample paths originate at the fundus of the lateral sulcus (Sylvian fissure) and end at area Spt, mid-post STS, and ATL, respectively. ***G***, Mean and standard deviation of activation phases centered at each selected node along the paths in ***F***.

### Circular statistics

The Watson-Williams test (Watson and Williams, 1956; Mardia and Jupp, 2009) and CircStat, a Matlab toolbox for circular statistics (Berens, 2009; https://www.mathworks.com/matlabcentral/fileexchange/10676-circular-statistics-toolbox-directional-statistics), were used to assess whether the activation phases at each node (vertex) on each path are significantly different between L1 and L2 across subjects. The complex *F*-values of *V* = 121 vertices (center and 120 neighbors) at node *d* on path *P* = {DVS, VVS, Spt, STS, ATL, DMS, VMS, AI} on the phase-encoded activation map of subject *S* = {1, 2, …, *N*} and language *L* = {L1, L2}, were averaged using:

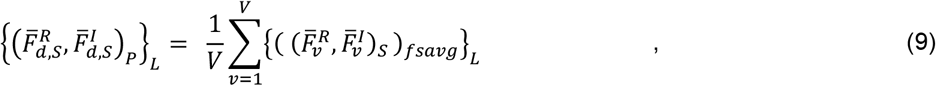

with a corresponding phase obtained by:

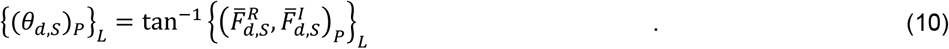

The distribution of phases across subjects at each node on each path is shown in Suppl. Figs. S1, S2, and S3. Given *N* complex *F*-values at node *d* on path *P* in L1 and L2 maps:

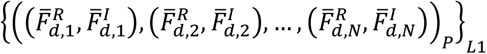

and 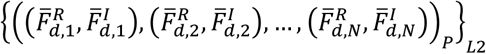, an *F*-statistic value of the Watson-Williams test is obtained by:

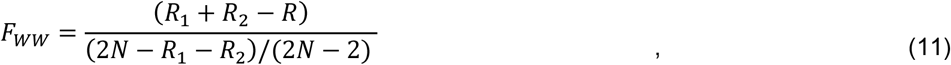

where *R*_1_, *R*_2_, and *R* are computed from two sets of phases, {(*θ*_*d,S*_)_*P*_}_*L*1_and {(*θ*_*d,S*_)_*P*_}_*L*2_, at each node *d* on each path *P* in *N* = 31 subjects using Equation 7.4.1 in Mardia and Jupp (2009). The resulting *F*_*WW*_ follows the *F*(1, 2*N*-2) distribution approximately (see *F*-values and *P*-values of all nodes on each path in Suppl. Dataset S1).

## Results

### Common and distinct cortical representations for L1 and L2 processing

The cortical representations of silent reading, listening, reading-aloud, and shadowing tasks in Chinese (L1) and English (L2) were mapped for each subject using rapid phase-encoded fMRI. For each task in L1 or L2, surface-based group average maps of statistically significant periodic activations (16 cycles per scan; *N* = 31; activation SNR: *F*(2, 230) > 4.7, *P* < 0.01; group statistics: *F*(2, 60) > 4.97, *P* < 0.01; cluster corrected) and their phases are color-coded and displayed on the inflated and flattened cortical surfaces of the template brain *fsaverage* (Fig. 1*A* to *D*). The commonalities and differences in activation extent between L1 and L2 are visualized in surface-based conjunction maps for each task (Fig. 1*E* to *H*). The group-average timings of recorded speech output during reading-aloud and shadowing tasks are summarized in Table 1.

For reading in L1 and L2 (Fig. 1*A, E*), activations were observed in the dorsal and ventral visual streams, middle to posterior portions of the superior temporal sulcus (mid-post STS), Sylvian parietal temporal area (Spt) at the posterior lateral sulcus, a strip of areas extending from the dorsal premotor cortex to the inferior frontal cortex (IFC), supplementary motor area (SMA), and pre-SMA in the left hemisphere. The right hemisphere showed no significant activation in areas Spt and mid-post STS for either language. Furthermore, reading in L2 activated a small area located at the junction of the frontal operculum (FOP) and anterior ventral insula (AVI) bilaterally, which was not activated during L1 reading.

At the same statistical significance level, reading in L2 consistently induced greater activation extent than L1 bilaterally in the inferior temporal gyrus (ITG), the dorsal visual stream along the intraparietal sulcus (IPS), dorsal and ventral premotor cortex, IFC, SMA, and pre-SMA (Fig. 1*E*). Reading in both languages activated smaller extent of the dorsal and ventral premotor cortex in the right hemisphere compared to the left. The conjunction map shows 98.7% of the vertices activated by L1 reading in the left hemisphere were also activated by L2 (Fig. 1*E*, Table 2). Conversely, only 57.6% of the vertices activated by L2 were shared with those activated by L1. For the right hemisphere, 96.9% of the vertices activated by L1 reading were also activated by L2, while just 51.9% of the L2-activated vertices overlapped with those from L1. This indicates substantial activation expansion from L1 to L2 in both hemispheres. Additionally, reading in L2 induced delayed activations (as indicated by bluish and greenish regions) in dorsal and ventral visual streams, dorsal and ventral premotor cortex, mid-post STS, area Spt, SMA, and pre-SMA in the left hemisphere (Fig. 1*A*).

Listening in both L1 and L2 activated the superior temporal cortex bilaterally (Fig. 1*B, F*), encompassing the superior temporal gyrus (STG) and extending into the mid-post STS. There was no significant difference in activation extent of this region between the L1 and L2 maps, as indicated by over 90% overlap of activations in both hemispheres (Table 2). The L2 map exhibited a slightly larger extent of activations in SMA, pre-SMA, area 55b in the dorsal premotor cortex, and IFC (including areas IFSa, IFSp, and IFJa according to the HCP-MMP1 atlas; Glasser et al., 2016), predominantly in the left hemisphere (Fig. 1*F*). Additionally, early visual areas activated in both L1 and L2 maps were engaged in processing a visual prompt (an ear icon) during listening tasks. Besides larger activation extent, listening in L2 also induced delayed activations in the temporal cortex, particularly in the mid-post STS and Spt (as indicated by greenish regions), along with SMA, pre-SMA, area 55b, and IFC in the left hemisphere (Fig. 1*B*). In contrast, the right hemisphere showed smaller activation extent in SMA and area 55b for both languages.

Compared with the regions activated during silent reading tasks (as indicated by solid contours in Fig. 1*C*), reading aloud in L1 and L2 engaged additional brain regions involved in language production (Fig. 1*G*). These include the middle to posterior portions of the cingulate sulcus and gyrus, dorsal and ventral premotor cortex, dorsal lateral prefrontal cortex (DLPFC), IFC, operculum, insula, secondary somatosensory cortex (SII), primary sensorimotor cortex (MI/SI; respiratory and orofacial representations), and parietal face and body areas (Sereno and Huang, 2006; Huang et al., 2012; Huang and Sereno, 2018). Beyond the regions commonly activated by reading aloud in both languages, L2 activated extended regions in DLPFC, IFC, superior and inferior parietal cortex, precuneus, anterior cingulate cortex, and ITG (see magenta regions in Fig. 1*G*). While reading aloud in L2 showed a greater expansion of activations in both hemispheres, only 62.3% of L2-activated vertices overlapped with those activated by L1 in the right hemisphere, slightly less than the 69.9% overlap in the left hemisphere (Table 2). Furthermore, reading aloud in L2 showed delayed activations relative to L1 in the premotor cortex, orofacial sensorimotor cortex, area Spt, mid-post STS, IPS, and ITG (as indicated by blue to greenish regions).

Shadowing in L1 and L2 activated bilateral superior temporal cortex and the same frontal regions engaged in reading aloud (Fig. 1*D, H*). The supplementary, premotor, primary sensorimotor, and insular regions involved in overt speech production were largely consistent between reading aloud and shadowing in each language (Fig. 1C, *D*). At the same statistical threshold, the extent of activation in the superior temporal cortex during shadowing was slightly greater than that during listening in the same language (as indicated by the contours in Fig. 1*D*). Furthermore, a protrusion in the anterior STS was activated during shadowing but not during listening in either language. The conjunction map for shadowing tasks indicated ∼80-90% overlap in activations between L1 and L2 (Fig. 1*H*; Table 2). In addition to comparing the extent of activations, we also found that shadowing in L2 exhibited delayed activations in the SMA, dorsal and ventral premotor cortex, and FOP/AVI (Fig. 1*D*).

The value in each unshaded cell represents the percentage of vertices that show significant activations (*P* < 0.01; cluster corrected) for either L1 or L2 alone, respectively represented by yellow and magenta regions in the conjunction maps (Fig. 1*E* to *H*). The value in each shaded cell represents the percentage of vertices with significant activations in both languages (overlap), indicated by the cyan regions.

### Language perception surfs through visual and auditory streams

Within the regions activated by L1 or L2 tasks (Fig. 1), the phase-encoded color patterns revealed hemodynamic traveling waves propagating across the cortical surface. As an initial step toward analyzing the complex spatiotemporal patterns, we sampled surface-based paths — analogous to extracting sample cores from tree rings — in the visual, auditory, and motor systems (Fig. 2*A*-*C*). We then compared the timings of traveling waves passing through each fixed path between L1 and L2 processing (e.g., Fig. 2*D,F*; Suppl. Dataset S1). All paths show progressive increases in activation phases from their starting points. The surface-based regions of interests that each path traverses are summarized in Table 3. Here, we report results with sROIs primarily according to the HCP-MMP1 parcellation (Glasser et al., 2016).

Reading tasks in L1 and L2 activated a dorsal visual stream (DVS) traversing the IPS and a ventral visual stream (VVS) traversing the ventral occipitotemporal cortex (Fig. 2*D*), with the L2 map exhibiting a greater extent of activations in both streams. Along the DVS path, L2 processing exhibited significant delays (*P* < 0.05, Watson-Williams test) in activation phases compared with L1 throughout the entire path (Fig. 2*E*, upper panel). The average delay for L2 processing in the DVS path was 0.97 s (Suppl. Dataset S1). Along the VVS path, L2 processing began to exhibit significant delays (*P* < 0.05, Watson-Williams test) compared with L1 at approximately one-third of the way along the path (Fig. 2*E*, lower panel).

Listening tasks in L1 and L2 induced a wide range of traveling waves originating at the fundus of the lateral sulcus and propagating across the superior temporal cortex through three major streams, ultimately reaching area Spt, mid-post STS, and anterior temporal lobe (ATL) (Fig. 2*F*). The sROIs along these paths are summarized in Table 3. In the first half of the Spt and STS paths, there was no difference in activation phases between L1 and L2. However, approximately halfway through these paths (Fig. 2*G*, upper and middle panels), L2 processing began to exhibit delayed activations compared with L1, with some nodes exhibiting significant delays (*P* < 0.05, Watson-Williams test). On the other hand, the entire ATL path showed no differences between L1 and L2 processing (Fig. 2*G*, lower panel).

### Language perception and production surf through multimodal streams

Reading aloud in both L1 and L2 induced complex traveling wave patterns in the visual, auditory, insular, supplementary motor, premotor, and primary sensorimotor cortices (Fig. 3). Our analysis tracked the progression of these waves along the same paths selected for reading and listening (Fig. 3*A, C*), paths DMS and VMS traversing the premotor and sensorimotor cortices (Fig. 3*E*), and path AI originating from the FOP/AVI (Fig. 3*E*). Along the DVS path traversing the posterior parietal cortex, L2 processing exhibited significant delays (*P* < 0.05, Watson-Williams test) in activation phases compared with L1 throughout this path (Fig. 3*B*, upper panel). On average, L2 lagged 0.88 s behind L1 processing along the DVS path (see Suppl. Dataset S1). Delays for L2 began around the midpoint and became statistically significant near the end along the VVS path traversing the ventral occipitotemporal cortex (Fig. 3*B*, lower panel). For the Spt path traversing the lateral sulcus, processing of L2 showed minor delays throughout the path, exhibiting statistical significance near area Spt (Fig. 3*D*, upper panel). About halfway through the STS path, L2 processing began to exhibit slight delays compared with L1 (Fig. 3*D*, middle panel). The entire ATL path displayed marginal delays for L2 compared with L1 (Fig. 3*D*, lower panel). In the DMS path, significant delays (*P* < 0.05, Watson-Williams test) were observed for L2 processing for most of the path (Fig. 3*F*, upper panel). Here, L2 was delayed by an average of 0.73 s compared with L1 (Suppl. Dataset S1). The VMS path also showed significant delays (0.94 s on average) for L2 processing, except in the last quarter of the path, where the waves arrived at the sensorimotor cortex (Fig. 3*F*, middle panel). The AI path exhibited slight delays for L2 processing most of the time during the first two-thirds of the path (Fig. 3*F*, lower panel).

**Figure 3.**
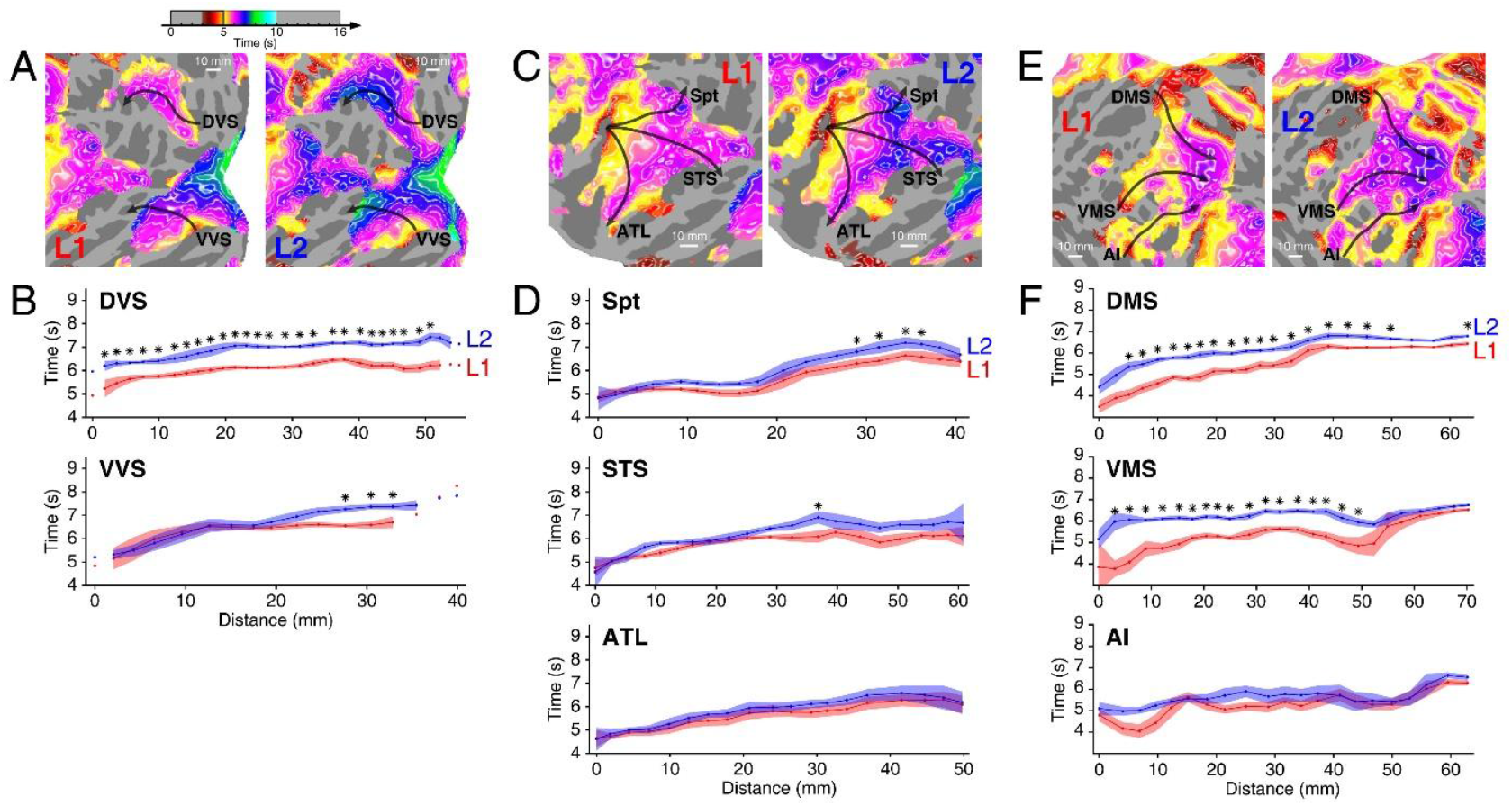
Analyses of processing times on multimodal streams for reading-aloud tasks in L1 and L2. ***A***, Closed-up views of traveling waves propagating through dorsal and ventral visual streams (see full-hemisphere maps in Fig. 1*C*). ***B***, Activation phases at selected nodes along the DVS and VVS paths shown in ***A. C***, Closed-up views of traveling waves propagating through three auditory streams (Spt, STS, and ATL) in the superior temporal cortex. ***D***, Activation phases at selected nodes along each path depicted in ***C. E***, Closed-up views of traveling waves propagating through streams in the dorsal and ventral motor system (DMS and VMS) and anterior insula (AI), reaching the primary sensorimotor cortex (MI/SI). ***F***, Activation phases at selected nodes along each path shown in ***E*.** All conventions are consistent with those in Fig. 2.

Shadowing in L1 and L2 induced traveling waves in the auditory, insular, supplementary motor, premotor, and primary sensorimotor cortices (Fig. 4*A, C*). Overall, the activation phases of traveling waves in these shadowing tasks were delayed relative to those during reading aloud. For the three auditory paths, the starting times for reading-aloud tasks in both languages ranged from 4.5 to 4.9 s, while in shadowing, they extended to between 5.5 and 6 s. Similarly, for the three motor paths, the starting times for reading-aloud tasks ranged from 3.5 to 5.1 s, while starting times for shadowing tasks spanned from 4.5 to 6.7 s (Suppl. Dataset S1). Consequently, all auditory and motor paths analyzed for shadowing (Fig. 4) showed overall temporal offsets compared with the corresponding paths analyzed for the reading-aloud tasks (Fig. 3). The offsets persisted throughout all three motor paths analyzed (Suppl. Dataset S1). Analysis of speech recordings confirmed slower average speech onset timings during shadowing than during reading aloud for the same language (Table 1). Among the three paths in the superior temporal cortex, Spt and STS paths exhibited insignificant delays in L2 processing in later sections, while the entire ATL path showed no differences between L1 and L2 processing (Fig. 4*B*). The three motor paths exhibited similar progression profiles between reading aloud and shadowing (Figs. 3*F*, 4*D*). Along the VMS path, shadowing in L2 exhibited significant delays (*P* < 0.05, Watson-Williams test) compared with L1 during the first two-thirds of the path (Fig. 4*D*, middle panel). However, there were no significant differences between shadowing in L1 and L2 across the entire DMS path (Fig. 4*D*, upper panel). The AI path exhibited slight delays for L2 processing for most of the path (Fig. 4*D*, lower panel).

**Figure 4.**
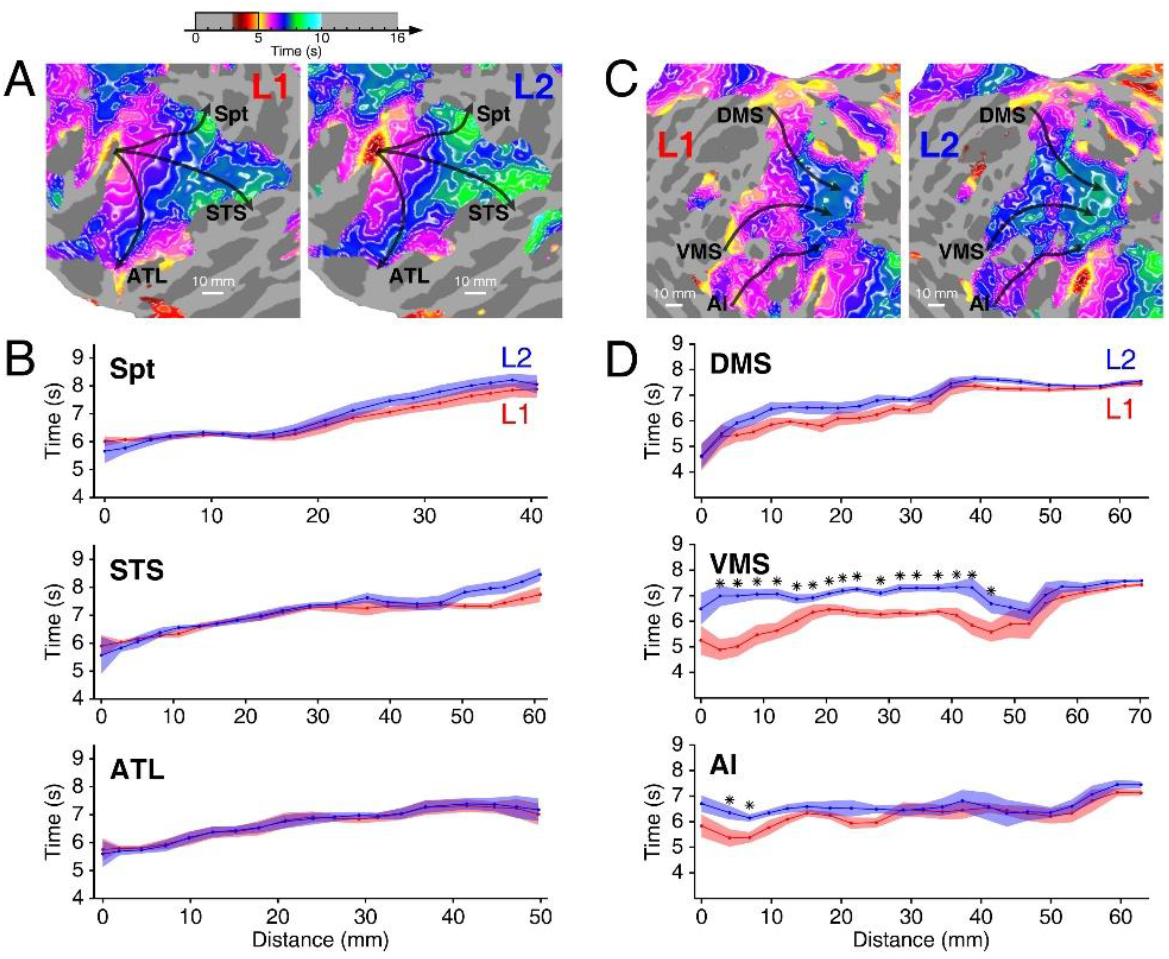
Analyses of processing times on multimodal streams during shadowing tasks in L1 and L2. ***A***, Closed-up views of traveling waves propagating through Spt, STS, and ATL streams in the superior temporal cortex during L1 and L2 shadowing tasks (see full-hemisphere maps in Fig. 1*D*). ***B***, Activation phases at selected nodes along each path illustrated in ***A. C***, Closed-up views of traveling waves propagating through DMS, VMS, and AI streams, reaching the primary sensorimotor cortex. ***D***, Activation phases at selected nodes along each path shown in ***C***. All conventions are consistent with those in Fig. 2.

## Discussion

Originally developed for topological mapping in sensory cortices (Engel, 2012), the rapid phase-encoded fMRI design with a short cycle (16 s) employed in this study successfully captured complex spatiotemporal activation patterns during naturalistic language perception and production tasks in sequential bilinguals. This method enables us to track hemodynamic traveling waves with comprehensive cortical coverage, revealing propagation patterns of brain activities aligned with findings from invasive techniques (e.g., Woolnough et al., 2019; Castellucci et al., 2022; Kovacs et al., 2025). Our results confirmed that L1 and L2 are processed through largely overlapping streams in visual, auditory, and motor systems. By analyzing the activation phases of these traveling waves along sample paths in each system, we identified specific locations and timings where L1 and L2 processing diverge. However, it is important to note that distinct micro-circuits for each language could yield similar activation phases, particularly in earlier sections of visual and auditory paths. While we define each one-dimensional path by selecting the starting and ending points of a hypothetical stream, the complex two-dimensional formations within each cortical region (e.g., the superior temporal cortex in Fig. 2*B*) warrant further spatiotemporal pattern analyses to achieve a full comparison of L1 and L2 processing. Building on this initial analysis of traveling waves, we will discuss the major findings below.

### Common and distinct activations between L1 and L2 processing

Conjunction maps revealed that silent reading and reading aloud in L2 exhibited larger activation extent in dorsal and ventral visual streams compared with L1 (Fig. 1*E, G*). In contrast, activation in the superior temporal cortex did not differ significantly between languages during listening, reading-aloud, and shadowing tasks (Fig. 1*F*-*H*). This might reflect fundamental differences in attentional control between L1 and L2. For instance, during L2 reading, subjects can voluntarily regulate their processing pace through eye movements, engaging a slower, more controlled process (Walczyk, 2000). In contrast, L2 listening relies on passive perceptual mechanisms within the auditory cortex, making it a relatively automatic process (Michael et al., 2001; De Santis et al., 2007). In addition to visual regions, the SMA, pre-SMA, and prefrontal cortex exhibited consistently greater activation extent during L2 tasks (Fig. 1*E*-*H*). These regions support the planning of preverbal articulatory movements (Indefrey and Levelt, 2004; Tourville and Guenther, 2011; Lima et al., 2016), suggesting that L2 invokes a greater cognitive demand in both perception and production.

While fMRI provides high spatial resolution for conjunction analysis between languages, it primarily reveals the strength and extent of hemodynamic activations, which limits our understanding of cognitive efficiency. Real-time language processing is inherently dynamic, requiring imaging both temporal and spatial brain activities (Hagoort, 2017). Although behavioral and EEG studies have demonstrated different temporal processing in L2 (Liu and Perfetti, 2003; Højen and Flege, 2005; Moreno and Kutas, 2005; Khateb et al., 2015; Perez et al., 2018), the precise neural origins of these differences remain unclear. By tracking the flows of hemodynamic traveling waves, we identified streams and locations where activation phases lag within overlapping visual, auditory, and motor regions during L2 language perception and production tasks (Figs. 2-4). The observed phase differences between L1 and L2 processing offer a direct temporal metric of cognitive efficiency.

### Temporal delays in visual streams during L2 reading

We observed that L2 processing lagged behind L1 in both silent reading and reading-aloud tasks across both visual streams (Fig. 2*D, E*; Fig. 3*A, B*). The DVS (the ‘where’ pathway) plays a crucial role in spatial awareness and sub-lexical analysis, such as directing eye movements during reading (Borowsky et al., 2006; Levy et al., 2009). The observed delay in DVS during L2 reading, coupled with a larger activation extent in the SMA, pre-SMA, posterior parietal cortex, premotor cortex, and IFG (Fig. 1*E, G*), indicates heightened attentional demand and a more controlled process required to parse a less familiar orthography. More critically, a significant delay was noted in the later sections of VVS (the ‘what’ pathway), which is responsible for automatic, lexical-level mapping of whole words to meaning (Borowsky et al., 2006; Levy et al., 2009). This slowdown in the VVS suggests that L2 reading encounters a bottleneck in accessing meaning, with this divergence localized to the fusiform gyrus (Figs. 2*D, E*; Fig. 3A, *B*).

### Temporal delays in auditory streams depend on speech sources

Our findings reveal a nuanced dissociation between L1 and L2 processing within the auditory dual-stream framework (Hickok and Poeppel, 2007; Saur et al., 2008; Rauschecker and Scott, 2009). For L2, processing delays emerged approximately halfway along auditory streams. The dorsal auditory stream (Spt), which is important for auditory-motor integration, showed significant delays in its later sections in L2 listening (Fig. 2*G*). During reading aloud in L2, however, most of the Spt path exhibited insignificant delays relative to L1 (Fig. 3*D*). This suggests that auditory-motor integration is less demanding for self-generated speech, as the internally predicted auditory signal requires less processing than an unpredictable external one.

In the ventral auditory stream, the STS path associated with phonological processing exhibited significant L2 delays in its mid-late sections during listening (Fig. 2*G*). However, the phase differences between L1 and L2 were markedly reduced during reading aloud (Fig. 3*D*). This is likely due to the predictability of self-generated auditory input, which lessens the demand for intensive phonological analysis. Conversely, the ATL path, related to semantic integration, showed no difference between L1 and L2 processing during both listening and reading aloud. We suggest that for competent bilinguals the simple S-V-O structure used in both languages did not necessitate deep semantic processing or judgement, resulting in no significant delay in L2 processing along the ATL path. Concurrently, the delays in the dorsal auditory-motor path (Spt) and the ventral phonological path (STS) are modulated by the speech source (external vs. self-generated), highlighting that the primary bottlenecks in L2 processing occur during the transformation of sound into articulation and phonological representations, with these bottlenecks being less pronounced for self-generated speech.

Two explanations are proposed for the absence of significant delays in all auditory streams during the shadowing task (Fig. 4*A, B*). First, the task’s immediate reproduction requirement may rely on a shallow acoustic encoding strategy, bypassing deeper phonological analysis. Second, the high cognitive demand of shadowing may lead to a ceiling effect, fully engaging the processing capacity of the auditory streams for both languages, thus eliminating the advantage of L1 processing. In summary, shadowing appears to saturate the typical language comprehension and auditory feedback networks, equalizing processing times between L1 and L2.

### Articulatory planning is affected by language nativeness during speech production

In the motor domain, significant delays in L2 processing during both reading aloud (Fig. 3*E, F*) and shadowing (Fig. 4*C, D*) along the VMS path suggest a consistent bottleneck in L2 articulatory planning. The ventral premotor cortex, which is critical for phoneme selection (Ghosh et al., 2008; Price, 2012; Planton et al., 2013; Hickok et al., 2023), appears to be less automated in L2. Notably, significant delays specific to the DMS path were observed only during reading-aloud, not during shadowing. The DMS involves regions, including area 55b, crucial for pitch coordination (Hickok et al., 2023). During shadowing, the external auditory input primes motor planning and output for both languages. In contrast, reading aloud requires the brain to generate the entire motor speech sequence from text internally. This internally driven generation and monitoring process places significantly more demand on the DMS, resulting in delayed activations. Lastly, the AI path, originating at the FOP/AVI junction, showed no significant L1-L2 differences (Figs. 3*F*, 4*D*). This path is distinct from the VMS path on the cortical surface. While its exact role in speech production remains uncertain, the anterior insula is thought to play a role in controlling respiration and auditory integration (Ackermann and Riecker, 2010; Oh et al., 2014; Woolnough et al., 2019). Importantly, respiration, as an autonomic function, is not influenced by the nativeness of a language.

### Conclusions

In this study, rapid phase-encoded fMRI introduced a temporal dimension to mapping spatiotemporal processing of languages in the bilingual brain. We demonstrated a methodological framework for analyzing the complex traveling wave formations of L1 and L2 processing through sample paths in visual, auditory, and motor systems. We identified precise temporal dissociation between L1 and L2 processing, and localized the delays in L2 processing to specific sections on shared paths. We proposed functional explanations for challenges associated with L2 processing. Specifically for silent reading and reading aloud in L2, the slower processing was due to heightened attentional demand through DVS and VVS; for L2 listening, the challenges lie in both phonological analysis and auditory-motor integration in the STS and Spt paths, respectively; for L2 production, bottlenecks in processing were identified in pitch coordination and articulatory planning along the DMS and VMS.

## Acknowledgment

This research was supported by the University of Macau Development Foundation (EXT-UMDF-014-2021); University of Macau (MYRG-CRG2024-00047-ICI, MYRG-GRG2023-00239-FAH, CPG2023-00016-FAH, MYRG2022-00265-ICI, MYRG2022-00200-FAH, CRG2021-00001-ICI, CRG2020-00001-ICI, SRG2019-00189-ICI, PIDDA 2020, PIDDA 2019); Macau Science and Technology Development Fund (FDCT 0001/2019/ASE); National Institutes of Health (R01 MH081990 to M.I.S and R.-S.H). We thank Yi Tang and Mengying Guo for help with fMRI experiments; Yafang Li, Aotong Li, and Ut Meng Lei for data preprocessing.

## Author Contributions

T.I.L, V.L.C.L., D.L., and R.S.H designed research; T.I.L, C.T.L., C.U.C., and R.S.H performed research; J.H.A and M.I.S contributed analytic tools; T.I.L., J.H.A, and R.S.H. analyzed data; T.I.L., V.L.C.L, and R.S.H wrote the paper.

## Code availability

Custom codes for analyzing phase-encoded fMRI data and traveling waves are included in csurf (a FreeSurfer-compatible package) available at https://pages.ucsd.edu/~msereno/csurf/

## Supplemental Material

**Fig. S1.**
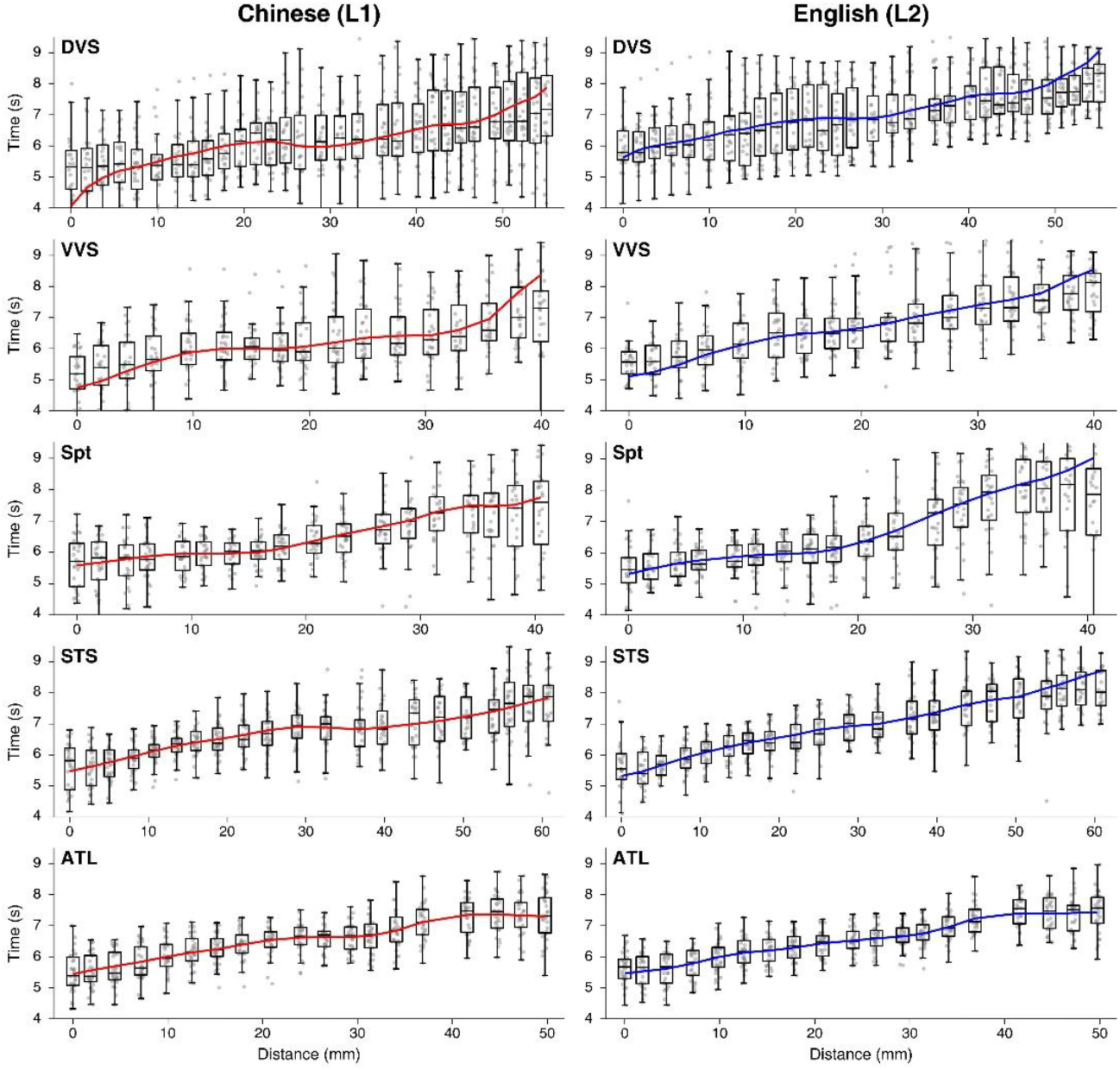
Analyses of L1 and L2 processing times on multimodal streams for reading and listening tasks in 31 subjects. Red and blue curves denote the corresponding group-average phase curves from Fig. 2. Each boxplot shows the distribution of single-subject activation phases at each node along the visual (DVS, VVS) and auditory (Spt, STS, ATL) paths.

**Fig. S2.**
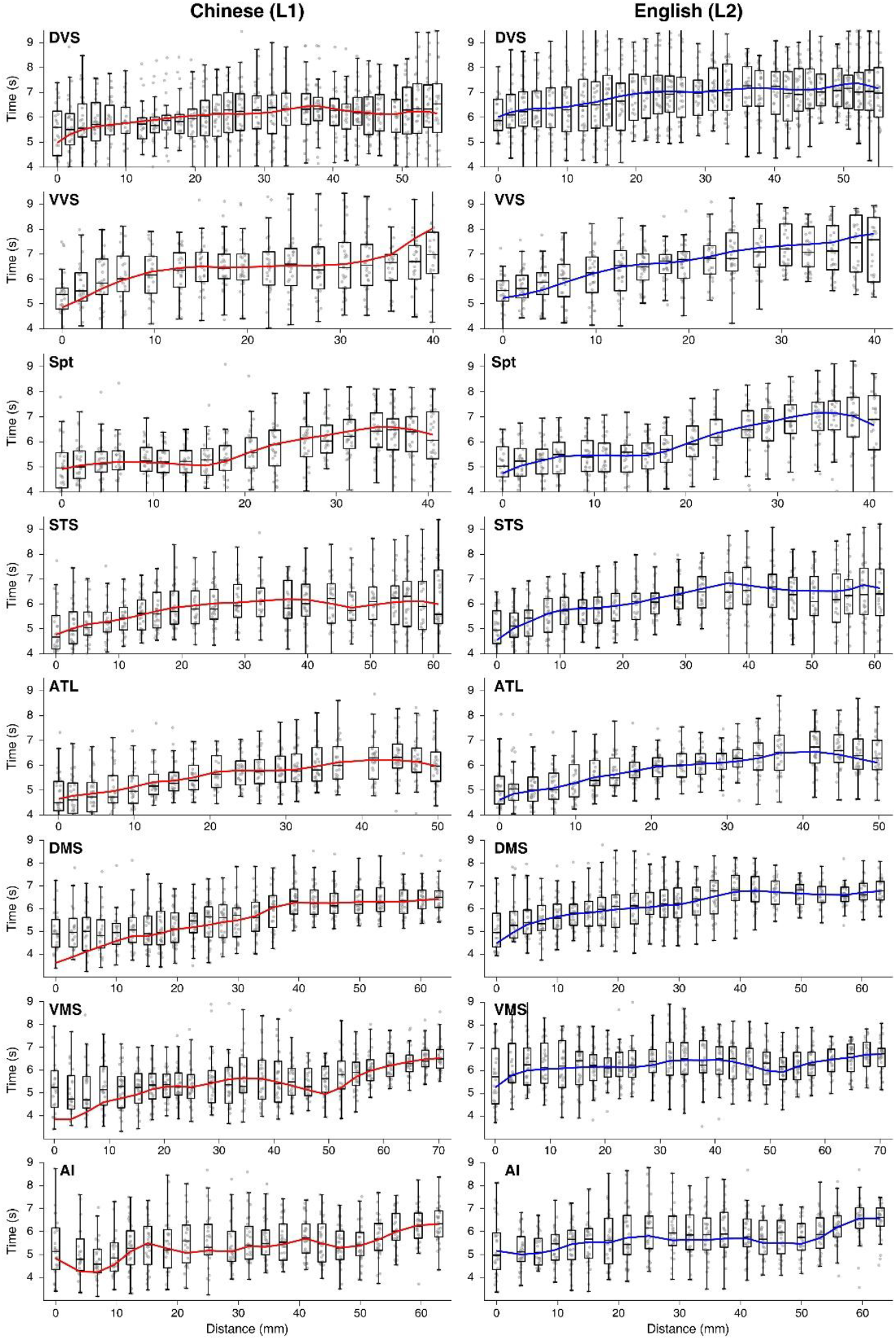
Analyses of L1 and L2 processing times on multimodal streams for reading-aloud tasks in 31 subjects. Red and blue curves denote the corresponding group-average phase curves from Fig. 3. Each boxplot shows the distribution of single-subject activation phases at each node along the visual (DVS, VVS), auditory (Spt, STS, ATL), and motor (DMS, VMS, AI) paths.

**Fig. S3.**
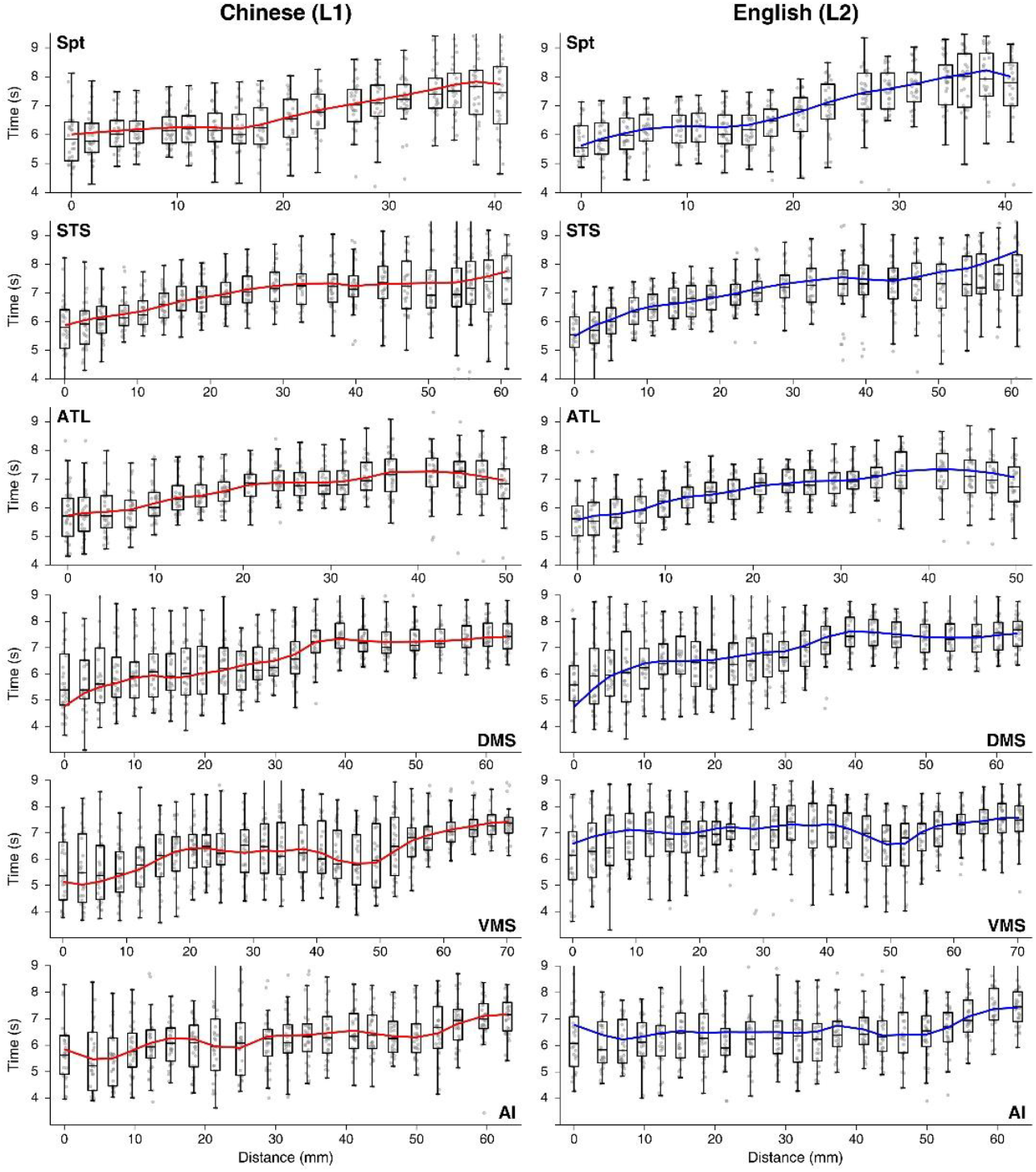
Analyses of L1 and L2 processing times on multimodal streams for shadowing tasks in 31 subjects. Red and blue curves denote the corresponding group-average phase curves from Fig. 4. Each boxplot shows the distribution of single-subject activation phases at each node along the auditory (Spt, STS, ATL) and motor (DMS, VMS, AI) paths.

### Dataset S1

Descriptive statistics for phase analyses in Figs. 2-4. Each Excel sheet lists the mean phase and standard deviation (s.d.) at each node on visual, auditory, or motor paths. Greyed-out values indicate unconnected nodes with insignificant activations (*P* > 0.01, cluster corrected) in the group-average maps. Each green-shaded cell indicates the mean phase of L2 processing lags behind L1 (i.e., L2 - L1 > 0). Each yellow-shaded cell indicates a significant difference between L1 and L2 phases tested across 31 subjects at each node (*F*(1,60); *P* < 0.05; Watson-Williams test).

